# NPZ*f*_*c*_: An ecological relation based fish catch prediction model using Artificial Neural Network

**DOI:** 10.1101/2023.11.08.566233

**Authors:** Manasi Mukherjee, Vettath Raghavan Suresh, Suman Kundu

## Abstract

Quantifying interactions of organisms of the various trophic levels is very important in understanding the dynamics of aquatic ecosystems. With regards to fish, as both ecologically and commercially important parts of an ecosystem, predicting their catch in relation to primary producers provides insight into sustainable management. The present paper describes a novel model NPZ*f*_*c*_, for nutrients, phytoplankton, zooplankton and fishes, which can predict planktivorous fish catch. Unlike the existing models which deal with the interactions within the system through mathematical equilibrium, the proposed model uses artificial neural network (ANN) to automatically learn inter-dependencies between different related variables and predict the fish catch of a water body. The efficiency of the model was increased by refining the input variables. Here biomass of plankton species population (phytoplankton and zooplankton) were specifically selected from feeding ecology studies of target fish species as input variable. The study of two of the commercially important fish species, *Etroplus suratensis* and *Nematalosa nasus* in Chilika lagoon showed that the model can predict with high accuracy from limited input data. The root mean square error (RMSE) is found to be very satisfactory, ranging from 3.53% to 11.5% for *E. suratensis* and from 1.63% to 2.22% for *N. nasus*. Higher accuracy and better predictive ability with a smaller dataset makes this ANN-based NPZ*f*_*c*_ model more conducive.

## Introduction

The increasing demand for natural resources and accounting their sustenance has constantly drawn attention to simulating the real world. Scientific models are widely used methods for substituting real-world systems into numerical relations. It allows experimenting with different inputs and analyzes how the end product is affected. As aquatic ecosystems account for the highest natural resources, ecosystem models have been persistently used for their management (Slobodkin, 1960; Odum and Odum, 2000). The models range from using the simplest parameters like nutrients and plankton (Franks, 2002) to complex organisms including humans (Wandersee et al., 2012). Models have also been proposed for understanding inter and intra interactions between abiotic and biotic components (Fulton, 2010; Rose et al., 2010). However, less effort has been taken to quantify this interactive understanding. The present study is one such attempt in which the primary producers and consumers are utilized to build a predictive fish catch model using Artificial Neural Network (ANN). To the best of our knowledge, unlike mathematical modelling, this is the first-ever use of ANN in the study of nutrient, phytoplankton and zooplankton (NPZ) interactions to predict forage fish catch.

Machine learning (ML) tools are known for their use in modelling patterns within data by automatically learning the parameters of the systems (Theodoridis and Koutroumbas, 1999; Pal and Mitra, 2004). Artificial Neural Network (ANN) is one such ML tools which tries to model biological neural networks (Yegnanarayana, 2009) and follows a simple principle of learning from examples without specifying any task-specific rules. As a result, ANN has been successfully used in systems where mathematical relations are hard to observe from the data (Kuo-lin et al., 1995). This motivated us to use ANN in the prediction of forage fish catch from NPZ values. The study was conducted in Asia’s largest coastal lagoon, Chilika, a Ramsar site. Fisheries of the lagoon share 71% of the economic value of the ecosystem (Kumar, 2003) and have international importance due to the exported value of around half a million USD in foreign exchange (Mohanty et al., 2008). The lagoon is well known for its biodiversity; wherein it was graded B on an ecosystem health report card using water quality, fisheries and biodiversity, indicating good water quality and good habitat condition of fishes (Pattnaik, 2013). The biodiversity component of the report card used bird richness, benthic diversity, dolphin abundance and phytoplankton diversity (Pattnaik, 2013). However, the nutrient flow through the food chain was not brought to the attention of those analyzing the system. Although Mohanty and Adhikary (2013) mentioned that interest in studying the trophic status of the lagoon and its relation to fisheries has been reported since the 1930s, the available literature do not show any advancements in study since Jhingran (1963). No attempts have been made so far to understand the complex behavior of this nutrient, plankton and fish relation of the lagoon and develop a model that elucidates the dynamics of the system with a predictive ability. Developing an ecological model of this nature can be advantageous in understanding dynamic changes in nature utilizing the basic characters, predicting changes with regard to both biotic and abiotic components and in turn assist in the management of the lagoon.

## Materials and methods

The model NPZ*f*_*c*_ approached the use of three variables, nutrients (N), microphytoplankton (P) and microzooplankton (Z) to predict catch of the planktivorous fish species (*f*_*c*_). As the final output (resultant), i.e. forage fish catch was measured in tonnes, and the input variables (biotic) were converted to biomass. In this context both the microphytoplankton and microzooplankton were represented in carbon biomass (mgC/l) estimated from species-specific biovolume. Microplankton species confirmed from feeding ecology analysis of the target fish species (Mukherjee et al., 2016, 2017) were exclusively used for calculating input variable (microphytoplankton and microzooplankton) biomass.

### Data collection

The study site, Chilika, is situated in Odisha, along the north east coast of India, lying between 19°28′ and 19°54′ North latitude and 85°6′ and 85°35′ East longitude. The lagoon has estuarine characteristics due to precipitation, the influx of freshwater distributaries of Mahanadi river system and seawater influx from the Bay of Bengal.

Simultaneous nutrient and microplankton samples were collected at monthly intervals using standard methods (Eaton et al., 2005). A 20 micron net was used to collect microplankton samples and their biomass (mgC/l) was estimated using biovolume .Biovolumes estimated based on geometric shapes were then converted to carbon in peta grams following Menden-Deuer and Lessard (2000). The measure followed for diatoms was

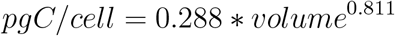

and for the rest protest plankton group the formula used was

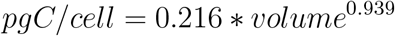

Planktivorous fish species catch data of the corresponding period have been obtained from Chilika Development Authority through a concurrent study.

### Data analysis

The data were non-linear. A predictive model for the data was attained by a supervised machine learning technique. In such learning, an algorithm called a classifier is trained with a set of labelled data to learn the relationship between the independent (nutrients, microphytoplankton and microzoo-plankton in this case) and the dependent variables (target fish species catch in this case). The machine learning technique used here was artificial neural network (ANN). In ANN multiple nodes (representing artificial neurons) are configured in layers with forward and optional backward communication links. The activity level of each node depends on the information received from the adjacent nodes of the previous layers. The learning nodes tend to send learned signals to the nodes in the next layer. Backpropagation of the results helps the neural network to reduce error. With continuous feeding of data, the classifier minimizes error to attain high accuracy in prediction. State of neurons is calculated as a weighted sum of received signals from neurons of the preceding layer. This is mathematically expressed as

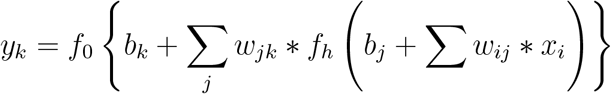

where, *y*_*k*_ = output signals, *x*_*i*_ = input signals, *w*_*ij*_ = weight between input neuron *i* to hidden neuron *j, b*_*j*_ = bias associated with hidden layer, *b*_*k*_ = bias associated with output layer, *f*_0_ = activation function for output layer and *f*_*h*_ = activation function for hidden layer.

The activation function of a neuron defines the output of that neuron based on weighted sum of the inputs given, i.e. it is a function that transforms the activation level of neuron to an output signal. Of the many activation function types (Bishop, 1995), the one used here for both the hidden and output layer is ‘tanh’. The tanh activation function is

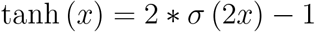

where, *σ* (*x*) is the sigmoid function. The range of tanh function is [-1,1] which provides stronger gradient. The error function used here with back-propagation was the least square error.

ANN used in the present study is configured with three input neurons representing the predictor variables nutrients, microphytoplankton and microzooplankton, one hidden layer with three neurons and one output neuron for the predicted dependent variable Fish (for e.g. *E. suratensis* and *N. nasus*). Similar to the constants in multiple regression, supplementary bias nodes were added to the hidden and output layer. A pictorial representation of the architecture is illustrated in Figure 1. On testing the performance with multiple hidden layers with different numbers of hidden nodes, only one hidden layer was selected due to its acceptable statistical performance. Equivalent results of a single hidden layer and multiple hidden layers have also been discussed often (Kůrková, 1992). The decision regarding the optimal number of nodes or neurons in the hidden layer was based on the performance comparison of varying networks. The number of hidden nodes in the network was tested varying from 1 to 5, and the number of nodes in the best performing network were chosen.

**Figure 1:**
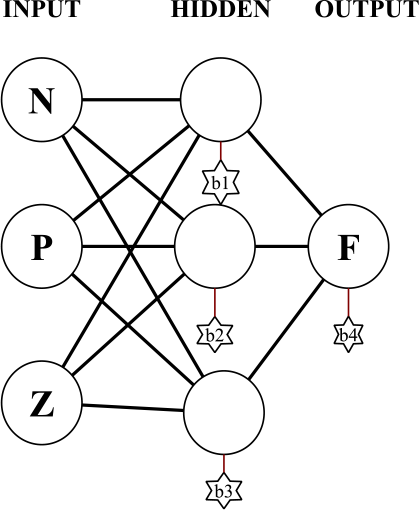
Neural network design of the model, with three input nodes and one hidden layer of three nodes. N = Nutrients, P = Microphytoplankton, Z = Microzooplankton, F = Fish. b1, b2, b3 and b4 are the biases obtained from each node.

Data set was split into training and testing samples where 80% were picked randomly for training and the rest were considered for testing. Ten such random training and testing samples were generated and used in ten different simulations. All of the training and testing values of the input [nutrients (here nitrate-N in ppm), microphytoplankton (units/l) and microzoo-plankton (number/l)] and output [fish catch (t)] were normalized using the logarithmic function. Output values of the test-set were then de-normalized before further statistical analysis. All of the machine learning algorithms were simulated and analyzed in the mathematical software ‘Mathematica 11’ (using inbuilt functions). The detailed steps of the process are given in Figure 2.

**Figure 2:**
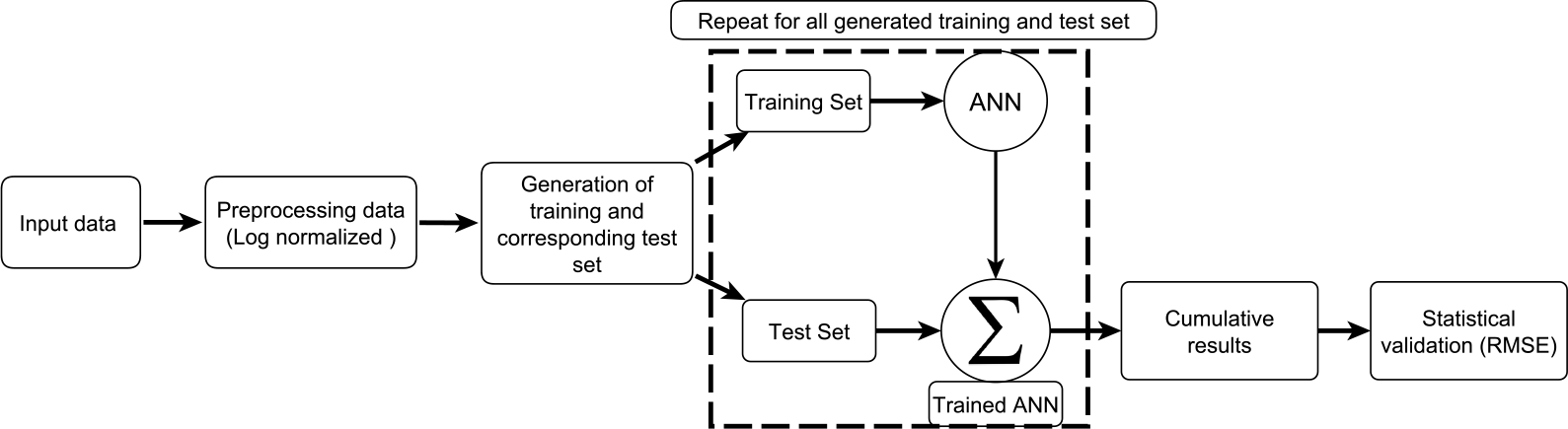
Steps followed in the NPZ*f*_*c*_ model

## Results

The model generated considered the transfer of energy at each trophic level in terms of biomass. Diet-specific plankton species of *N. nasus* and *E. suratensis* were selected from Mukherjee et al. (2016). The biovolume of these plankton species (Mukherjee and Suresh, 2019) was converted to biomass. The total microphytoplankton carbon biomass was calculated as 2.96 × 10^−3^ mgC/cell and of microzooplankton was calculated as 1.28 × 10^−3^ mgC/cell. The monthly average microphytoplankton abundance of the species measured for biovolume ranged from 20 units/l in December to 19711 units/l in April and microzooplankton ranged between 1 no./l in October and 1992 no./l during May (Table 1).

**Table 1:**
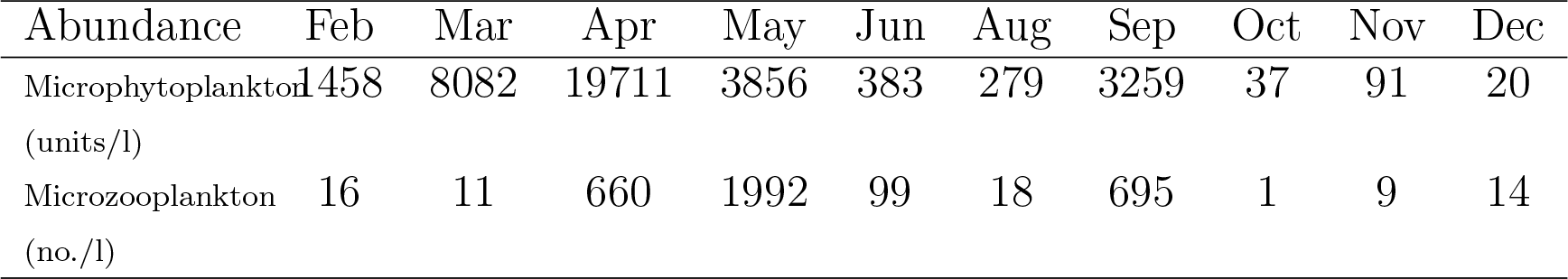
Monthly average abundance of plankton enumerated for biovolume

The biomass trend showed by microzooplankton followed that of micro-phytoplankton, wherein the gradual increase in phytoplankton from February to April was followed by an increase in zooplankton from March to May (Figure 3). The transfer of this biomass to the next trophic level was studied through the feeding ecology of two important forage fishes of Chilika viz. *E. suratensis* (Mukherjee et al., 2017) and *N. nasus* (Mukherjee et al., 2016). *E. suratensis* catch gradually increased and reached its peak in August when both the plankton groups showed a drop in biomass (Figure 3). A Further decline in the fish catch of September showed a corresponding increase in biomass of both groups of plankton. The catch trend of *N. nasus* also showed a similar relation to that of *E. suratensis*, wherein its catch increased with a decrease in plankton abundance (Figure 3). It is only during the month of April that *N. nasus* catch increased with microphytoplankton and a corresponding decrease in microzooplankton abundance. This fish species was found to have a very specific need for microzooplankton during the month which is discussed in details by Mukherjee et al. (2016). Thus the possibility of plankton growth based on the available nutrients of the environment and corresponding growth of planktivorous fish using them as food can be established for both species.

**Figure 3:**
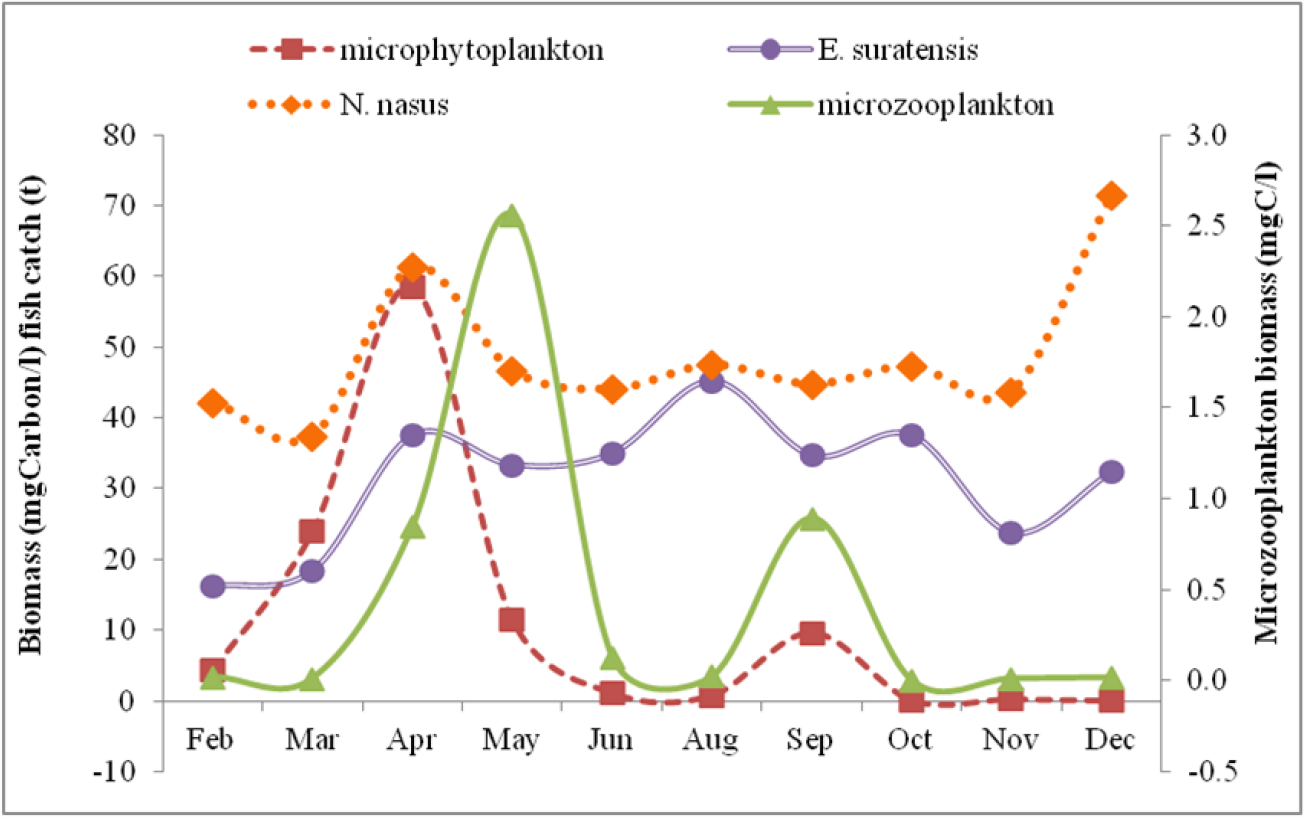
Monthly variation in biomass of plankton, *E. suratensis* and *N. nasus*

Table 2 reports the values of predictor or input variables of the model used along with the fish catch. It comprised of 10 months of sampling data with nitrate (mg/l) as nutrient, microphytoplankton (mgC/l), microzooplankton (mgC/l) and catch (t) of planktivorous fishes *E. suratensis* and *N. nasus*. The predicted catches (i.e., the output from the model) with the corresponding actual catches of all of the test samples are given in Figures 4a and 4b for *E. suratensis* and *N. nasus* respectively. Corresponding numerical values are mentioned in Tables 3 and 4.

**Table 2:**
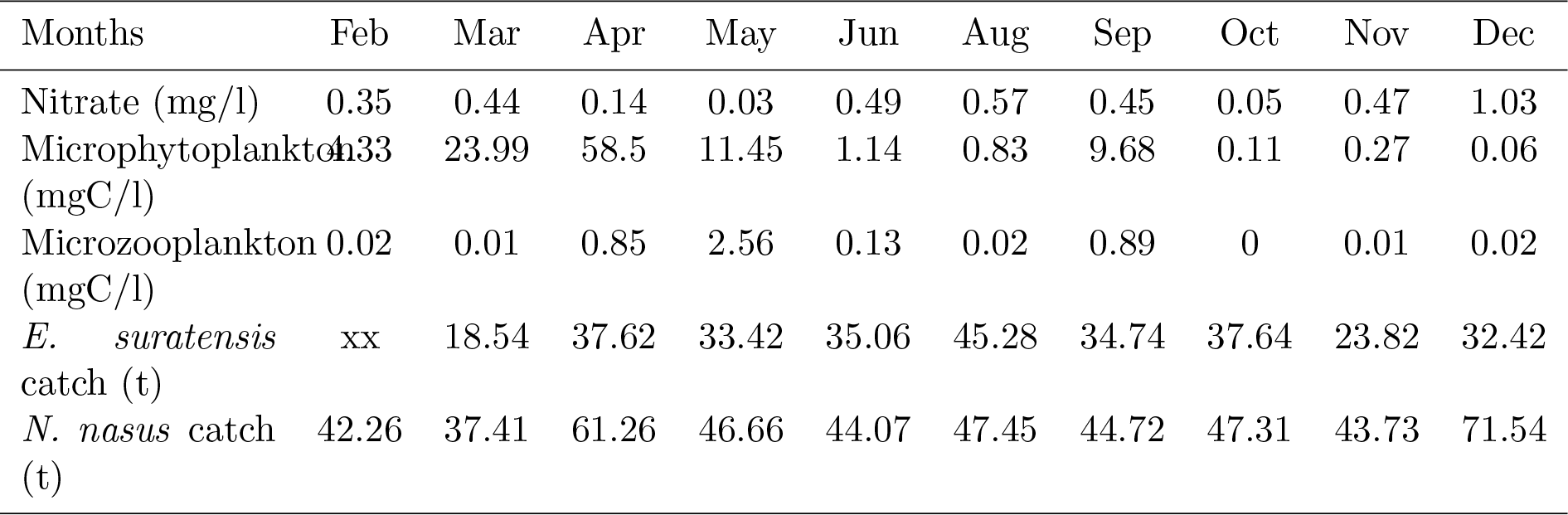
Input data of *E. suratensis* used for training and testing the model

| Months | Feb | Mar | Apr | May | Jun | Aug | Sep | Oct | Nov | Dec |
| --- | --- | --- | --- | --- | --- | --- | --- | --- | --- | --- |
| Nitrate (mg/l) | 0.35 | 0.44 | 0.14 | 0.03 | 0.49 | 0.57 | 0.45 | 0.05 | 0.47 | 1.03 |
| Microphytoplankton (mgC/l) | 4.33 | 23.99 | 58.5 | 11.45 | 1.14 | 0.83 | 9.68 | 0.11 | 0.27 | 0.06 |
| Microzooplankton (mgC/l) | 0.02 | 0.01 | 0.85 | 2.56 | 0.13 | 0.02 | 0.89 | 0 | 0.01 | 0.02 |
| *E. suratensis* catch (t) | xx | 18.54 | 37.62 | 33.42 | 35.06 | 45.28 | 34.74 | 37.64 | 23.82 | 32.42 |
| *N. nasus* catch (t) | 42.26 | 37.41 | 61.26 | 46.66 | 44.07 | 47.45 | 44.72 | 47.31 | 43.73 | 71.54 |

**Table 3:** Goodness of fit of model assessed through statistical tests

| Test Sample | Predicted | Actual | RMSE | %RMSE | MAE | %MAE |
| --- | --- | --- | --- | --- | --- | --- |
| 1 | 44.93 | 33.4 | 18.3 | 3.53 | 0.02 | 1.93 |
| 2 | 33.81 | 34.7 | 18.6 | 3.64 | 0 | 0.17 |
| 3 | 21.37 | 45.3 | 19.1 | 3.68 | 0.05 | 4.64 |
| 4 | 42.17 | 34.7 | 18.7 | 3.64 | 0.02 | 1.5 |
| 5 | 59.68 | 32.4 | 19.2 | 3.76 | 0.06 | 6.03 |
| 6 | 15.19 | 18.5 | 18.6 | 4.08 | 0.01 | 0.85 |
| 7 | 62.78 | 32.4 | 19.2 | 3.97 | 0.08 | 8.04 |
| 8 | 19.20 | 33.4 | 18 | 4.45 | 0.05 | 4.52 |
| 9 | 49.14 | 35.1 | 18.3 | 4.4 | 0.05 | 4.76 |
| 10 | 25.84 | 34.7 | 18.7 | 4.87 | 0.04 | 3.61 |
| 11 | 16.11 | 18.5 | 19.4 | 4.99 | 0.01 | 1.1 |
| 12 | 63.48 | 35.1 | 20.4 | 4.89 | 0.14 | 13.9 |
| 13 | 8.00 | 37.6 | 19.2 | 6.39 | 0.21 | 21 |
| 14 | 34.43 | 35.1 | 17.2 | 6.02 | 0 | 0.48 |
| 15 | 6.21 | 37.6 | 18.5 | 7.11 | 0.32 | 31.9 |
| 16 | 17.76 | 37.6 | 14.7 | 6.5 | 0.22 | 21.5 |
| 17 | 20.09 | 18.5 | 13 | 6.71 | 0.02 | 2.08 |
| 18 | 14.89 | 37.6 | 15 | 7.35 | 0.42 | 41.8 |
| 19 | 22.11 | 34.7 | 8.97 | 7.59 | 0.32 | 32 |
| 20 | 17.41 | 18.5 | 1.13 | 11.5 | 0.06 | 6.49 |

RMSE = Root Mean Square Error
MAE = Mean Absolute Error

**Table 4:** Goodness of fit *N. nasus* model assessed through statistical tests

| Test Sample | Predicted | Actual | RMSE | %RMSE | MAE | %MAE |
| --- | --- | --- | --- | --- | --- | --- |
| 1 | 41.303 | 61.260 | 11.137 | 2.020 | 0.020 | 2.015 |
| 2 | 45.891 | 47.450 | 10.469 | 2.002 | 0.002 | 0.164 |
| 3 | 44.920 | 43.730 | 10.750 | 1.993 | 0.001 | 0.132 |
| 4 | 46.569 | 44.720 | 11.058 | 1.981 | 0.002 | 0.215 |
| 5 | 42.978 | 37.410 | 11.389 | 1.972 | 0.007 | 0.686 |
| 6 | 45.441 | 43.730 | 11.674 | 1.952 | 0.002 | 0.223 |
| 7 | 40.550 | 42.260 | 12.075 | 1.936 | 0.002 | 0.237 |
| 8 | 42.452 | 47.310 | 12.522 | 1.905 | 0.007 | 0.712 |
| 9 | 46.020 | 44.070 | 12.958 | 1.875 | 0.003 | 0.305 |
| 10 | 40.323 | 42.260 | 13.521 | 1.852 | 0.003 | 0.326 |
| 11 | 46.463 | 47.450 | 14.168 | 1.806 | 0.002 | 0.178 |
| 12 | 65.772 | 37.410 | 14.931 | 1.774 | 0.056 | 5.590 |
| 13 | 70.860 | 44.720 | 12.257 | 1.812 | 0.059 | 5.920 |
| 14 | 48.438 | 47.450 | 8.608 | 1.888 | 0.003 | 0.267 |
| 15 | 45.436 | 47.450 | 9.288 | 1.862 | 0.006 | 0.625 |
| 16 | 47.144 | 46.660 | 10.135 | 1.806 | 0.002 | 0.175 |
| 17 | 45.441 | 43.730 | 11.329 | 1.742 | 0.007 | 0.745 |
| 18 | 69.950 | 71.540 | 13.044 | 1.628 | 0.009 | 0.863 |
| 19 | 69.194 | 46.660 | 15.936 | 1.750 | 0.197 | 19.717 |
| 20 | 45.090 | 44.720 | 0.370 | 2.218 | 0.008 | 0.820 |

RMSE = Root Mean Square Error
MAE = Mean Absolute Error

**Figure 4:**
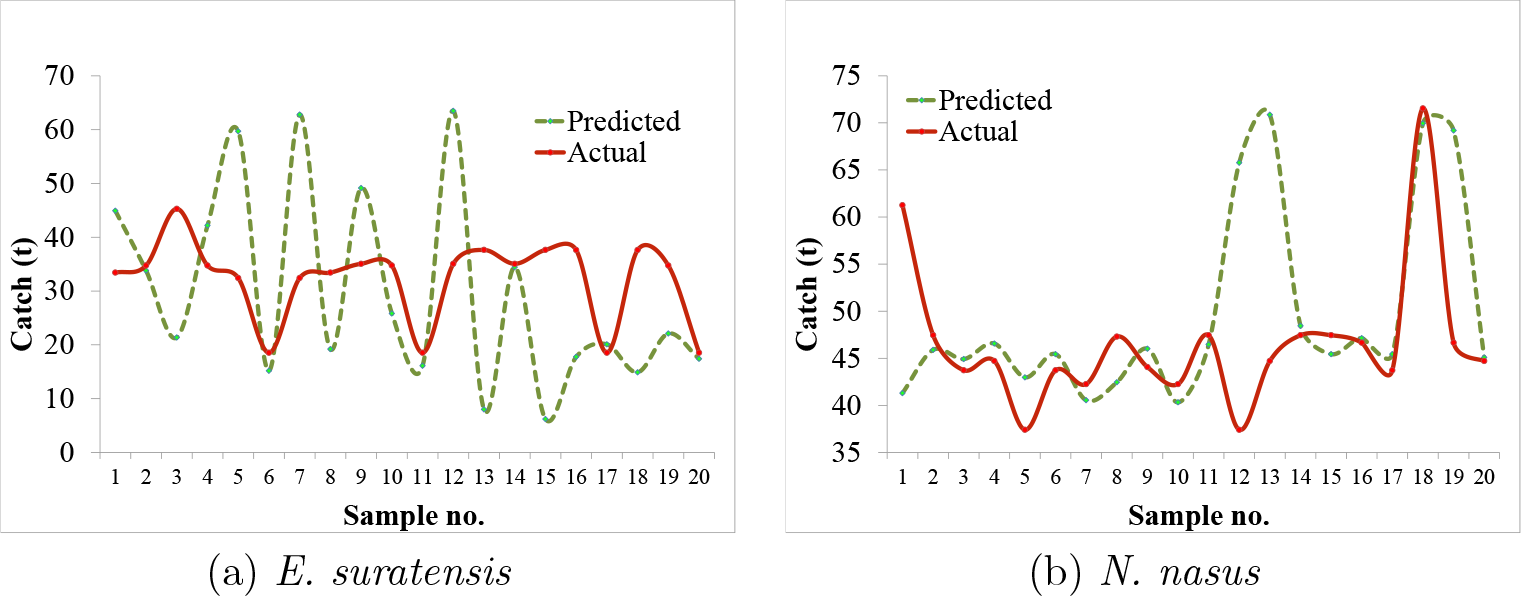
Predicted and actual catch of fish generated from 20 test set used for validation of the model

In the case of *E. suratensis*, 13 values (65%) remained higher and 7 lower (35%) predictions. Of these 65% higher predicted values, three values were almost perfect predictions, (0.93t, 0.63t, 1.13t) remaining within a difference of 1 t. For *N. nasus*, the higher and lower predicted values remained equal in percentage (50%), of which four values were near perfect (1.56t, 1.71t, -1.71t and 1.93t). The values for all 20 samples are given in Table 3 for *E. suratensis* and Table 4 for *N. nasus* for reference.

Root mean square errors (RMSEs) were calculated to find the variations amongst the actual and predicted value. Root mean square errors are more sensitive to occasional very large errors in predicted values as they tend to square the errors in the calculations. The mean absolute error (MAE) was also used to complement the idea of goodness of fit. Error values were also converted into percentages to better visualize the differences. The RMSE values of *E. suratensis* ranged from 1.13 t (3.53%) to 20.4 t (11.5%), while the mean RMSE was 16.7 t (5.45%) (Table 3). MAE calculated ranged from 0 t (0.17%) to 0.42 t (41.48%), with an average of 0.1 t (10.4%). The RMSE value of *N. nasus* ranged from 0.37 t (1.63%) to 14.93 t (2.22%), with the mean being 11.38 t (1.89%) (Table 4). MAE ranged from 0 t (0.13%) to 0.19 t (19.71%) with an average of 0.02 t (1.99%).

## Discussion

The production of an ecosystem depends upon the energy flow through each trophic level. Therefore, acquiring an idea regarding the fish production as an end product would be dependent on their source of energy. Chilika lagoon inhabits huge biodiversity and thus a complex trophic chain. Plankton dynamics of the ecosystem also has distinct spatio-temporal variations (Srichandan et al., 2015; Mukherjee et al., 2018). These variations have a considerable effect on the forage fish population, which in turn draws the attention of ecologists to develop compartment models (Kumar and Kumari, 2015; Franks, 2002). The present study finds the scope of establishing a link between plankton dynamics and feeding ecology studies to develop a fish catch model. A detailed study of feeding ecology of the *N. nasus* and *E. suratensis* (Mukherjee et al., 2016, 2017) has established them as planktivorous fishes. The relationship was further used to build a model that utilizes the transfer of energy from nutrients to microphytoplankton thereon to microzooplankton and finally to planktivorous fish. The relation between these four compartments provided a platform to build a model that can be predictive in nature. The four compartments here were developed into an NPZ*f*_*c*_ model, through the artificial neural network, that can be very useful over the traditional mathematical NPZ models (Franks, 2002) or recently developed NPZF models (Kumar and Kumari, 2015). The mathematical NPZ models are an effective approach to predict the dynamics in aquatic systems that are hard to measure for (Franks, 2002). However, the present work was interested in predicting or perhaps quantifying the final output based on the related inputs; an approach that sounds more statistical, as conventional methods like regression do the same. While various species of forage fishes are confirmed as specialized, preferential and generalized feeders(Mukherjee et al., 2016, 2017), considering the transfer of energy from the entire plankton community to the forage fishes in developing a model would be inappropriate. Thus, of the 233 plankton species recorded (Mukherjee et al., 2018), the present model uses the biomass of only 85 species of plankton that occurred in the target fish species’ diet. This approach of species specificity in plankton increases the accuracy of forage fish species catch prediction. The model can be very useful to predict catches of commercially important forage fish species of an ecosystem, using extensively generated feeding ecology data. This was addressed based on not mere statistical approaches but by more progressive learning to understand the complex inclusive variations in the environment. A number of statistical methods and tools are available that could be used to approach this model, but the requirement for a larger dataset was one of the most important constraints to do so. Moreover, the less the dataset fits into the conventional statistical tool (regression), the greater the chances of error and thus the lower the efficacy in prediction. As the collection of biological, specific environmental data remains difficult in terms of time and cost; large data sets are seldom available. Thus, an approach that could use smaller datasets and yet produce more effective predictions with fewer errors was needed. Machine learning was identified as one such potential tool that provides better results with iterative searches. Working through ANN provided the advantage of attaining good results relatively with such constraints, identifying the factors for change and summarizing the behavior of complex systems with small observational datasets (Pasini, 2015). The use of neural networks in phytoplankton production (Scardi, 1996; Mattei et al., 2018), succession (Olden, 2000) and bloom (Kang et al., 2012) predictions indicated the ability of these methods to work at the primary trophic level of an ecosystem. Thus the input and output data used for the generation of the current model were conducive in terms of biomass of primary producers (Table 2), signifying the transfer of energy in each trophic level. The use of ANN to relate environmental variables and fish catches (Iglesias et al., 2004; Gutiérrez-Estrada et al., 2009) established the networks efficacy all along the trophic levels of an ecosystem. The relationship between plankton abundance and fish catch has also been often discussed (Steingrund and Gaard, 2005) and approached for model development. These approaches were broad in terms of both the input and output variables, as far as the entire plankton population and pelagic fish population were regarded. As the variables here are related in terms of prey and predator, knowledge and use of specific food groups and fish species are more appropriate as variables and in turn would enhance the accuracy. In addition, concern of availability of large data sets can also be addressed with the same approach. *E. suratensis* and *N. nasus* are two of the most commercially important fish species of Chilika. As it is not migratory, the lake resident *E. suratensis* is better related to the corresponding changes in environmental variables (Figure 3). *N. nasus*, although migratory in nature, has very specific food utility from the environment (Mukherjee et al., 2016), providing a good relation with the variables (Figure 3). Thus, the study indicated that a predictive model based on such parameters requires a detailed catch structure of the target fish species to be studied and a probable residential species of an ecosystem to be considered.

The model showed good fit with very small data feeds for both of the fish species. Figure 4a shows that the predicted catch values of *E. suratensis* are on a par with the actual catch values for sample no.’s 1, 2, 4, 6, 10, 11, 14, 17 and 20, comprising 45% of the test data set. Perhaps the sample no. 6, 11, 14, 17 and 20 had exact match, which comprised 25% of the data. Of the remaining 55% of the data set, the data points 5, 7, 9 and 12 (20%) of predicted values remained larger than the actual values and the data points at 3, 8, 13, 15, 16, 18 and 19 remained lower. These resulted to a mean RMSE value of 11.38, i.e., only 5.45%. The model for *N. nasus* also was studied for differences in their predicted and actual catch (Figure 4b) and 55% were found to be on a par, i.e., about 1 t of difference. The mean RMSE for *N. nasus* was found to be even lower when compared to *E. suratensis*, as the value was only 1.89%. Thus, the proposed ANN-based model showed a good fit for both of the fish species studied and has an acceptable RMSE, indicating that the relationship established between nutrients, microphytoplankton, microzooplankton and planktivorous fish was appropriate to develop a predictive model of planktivorous fish catch. This NPZ*f*_*c*_ model, developed using ANN, was found to be an alternative approach to conventional mathematical modelling that is often used.

An ecosystem that supports the livelihood of more than 0.2 million (Pattnaik, 2013), shows that fish catch can be a very important tool for the practice of sustainable fishery. Fish catch predictions would therefore be an important tool towards ecological, ecosystem and socio-economic management. Planktivorous fish become food of the next trophic level, like carnivorous fishes. Catch of these carnivorous fishes can also be predicted by analyzing the prey species and their corresponding catch and fitting into the model. This model can thus be modified for any such species-specific or holistic approach of predicting related fish species, as the input variables have been analyzed in prior studies. The model is not only restricted to quantifying fishes, but can also be used in predicting any such environmental process that has definite corresponding related parameters measured. This study therefore signifies the importance of machine learning, especially ANN in developing ecological models that can be aimed at predicting results, measuring the magnitude of changes, and revealing important biological relationships by automated learning, thus having the potential to become an important tool in fisheries management.

## Acknowledgments

The authors are thankful to the Chilika Development Authority for providing the fish catch data of the targeted species. We are also grateful to Dr. R. K. Manna for aiding the nutrient data. Suman Kundu like to acknowledge the National Science Centre, Poland for the grant 2016/23/B/ST6/01735.

## Notes

### Competing Interest Statement

The authors have declared no competing interest.

